# Hidden components of microbiome diversity revealed by cell wall lysis with Adaptive Focused Acoustics

**DOI:** 10.1101/2020.05.21.109736

**Authors:** Giselle F. Wallace, Sonia Ahluwalia, Vishal Thovarai, James Laugharn, Hamid Khoja, Shurjo K. Sen

## Abstract

Within the rapidly evolving field of microbiome sequencing, a primary need exists for experimentally capturing microbiota in a manner as close as possible to their *in vivo* composition. During microbiome profiling, the first step necessarily involves lysis of the cell wall, releasing nucleic acids for next-generation sequencing. Microbial cell wall thicknesses can vary between 5nm to 80nm; while some species are quite easy to lyse, others are particularly resistant to lysis. Despite this, current chemical/mechanical lysis protocols ignore the possibility that species with different cell wall thicknesses are lysed at differential rates. This creates noise in species compositions and possibly skews current microbiome results in ways that are not currently understood. To develop a cell wall thickness-agnostic lysis protocol, we used Adaptive Focused Acoustics (AFA), a tunable acoustic methodology for processing of biological samples. Using identical aliquots of mouse stool homogenate as the lysis substrate, we compared AFA with chemical/mechanical lysis methodology routinely used in microbiome studies and found that AFA-mediated lysis substantially increases both microbial DNA yield as well as alpha and beta diversity. By starting with lower AFA energy levels, sequentially removing aliquots at each step, and subjecting the remainder to progressively stronger AFA treatment, we developed a sequential lysis method that accounts for differences in cell wall thickness. This method revealed even greater levels of diversity than single-timepoint AFA treatment. 16S sequencing results from the above experiments were verified by shotgun metagenome sequencing of a subset of the AFA samples. We found that lysis-induced noise affects not just species compositaions, but also functional characterization of shotgun metagenome data. AFA samples also showed a higher detection of eukaryotic and fungal DNA. We suggest that AFA-mediated lysis produces a truer representation of the native microbiota, and that this method deserves consideration as a potential addition to microbiome lysis protocols.

## Introduction

In recent years, knowledge of the community structure and functional composition of the microbiome and its role in human health has expanded dramatically.^1, 2^ However, concerns regarding a lack of reproducibility continue to restrict progress in microbiome research.^3–5^ In general, there remains an urgent need for consistent and broadly agreed-upon methods of microbiome sample processing, along with a better understanding of sources contributing to non-biological noise in microbiome sequencing data.^6, 7^ Sources of non-biological variation in current microbiome protocols include (but are not limited to) sample collection method, PCR amplification bias, next-generation sequencing (NGS) platform, and bioinformatics analysis technique.^8^ In particular, DNA extraction may be the stage at which a substantial amount of variability is introduced during microbiome sequencing.^9–11^ Here, we focus on bacterial cell wall lysis, the first step of DNA extraction, as a major area for exploration and improvement of reproducibility.^12, 13^

Currently, most bacterial lysis protocols involve mechanical bead-beating and chemical/enzymatic treatment, either singly or in combination.^14, 15^ However, despite our long-standing knowledge that bacterial cells walls possess a broad range of thicknesses and biochemical structures, existing protocols ignore the possibility that a one-size-fits-all cell wall lysis step may produce a skewed representation of native microbial community structure.^16^ Specifically, we suggest that delivery of a fixed amount of mechanical energy from traditional bead-beating lyses the cell walls of different taxa and releases DNA at rates that reflect their susceptibility to lysis rather than their native abundance in a microbial community. Hence, it is possible that current microbiome protocols are deficient in experimentally representing microbial diversity into NGS-derived taxonomic ratios (through over- or under-lysis of bacterial taxa at the lower and upper ends of the cell wall thickness spectrum, respectively).

Here, to address lysis-induced noise in microbiome sequencing, we carried out a systematic study of bacterial lysis techniques, incorporating both traditional chemical/mechanical protocols along with Adaptive Focused Acoustics (AFA),^17, 18^ a precise and scalable system that uses non-contact delivery of focused short-wavelength ultrasonic energy to fracture the cell wall.^19^ More specifically, AFA employs focused bursts of high-frequency ultrasonic acoustic energy (wavelength 1-3 mm) focused into a discrete zone within a sample vessel held at near-isothermal conditions (thus enabling cell wall lysis without heat-induced damage of nucleic acids). Although AFA has found widespread use in other NGS protocols,^20–22^ our study represents the first evaluation for use in microbiome sequencing. In addition to the side-by-side comparison of AFA with traditional chemical/mechanical protocols (where a single lysis step was used for both methods), we also tested the efficacy of a sequential AFA-based microbiome lysis technique, which aims to account for variation in bacterial cell wall thickness and rigidity in a microbiome sample. For this protocol, sub-aliquots of the same starting sample were exposed to varying levels of acoustic energy and then combined prior to sequencing. From our experiments, we found that when identical stool homogenate aliquots were processed using AFA-based lysis and bead beating, AFA processing yielded more microbial DNA and revealed greater levels of both alpha and beta diversity. Further, we found that numerous bacterial taxa were detected with greater success in AFA-lysed samples relative to bead-beating, and that higher levels of eukaryotic diversity were found by this lysis method. We show that lysis-associated taxonomic differences affect functional interpretation of microbiome data. In general, our results suggest that a certain component of species diversity in microbiome samples may be missed by existing lysis protocols. Our evaluations of AFA-based lysis methods are hoped to serve as a resource to the community in finding the best possible solution for microbiome workflows, i.e. one that will be effective, fast, reproducible, and also account for differences in lysis susceptibility for the diverse communities of species in microbiome samples.^23^

## Materials and Methods

### Mouse stool collection, cell wall lysis and DNA extraction

Stool pellets from C57BL/6 mice were collected over dry ice and pooled together in a 50ml Falcon tube. Post-collection, 5 mL of TE buffer pH 8.0 (Amresco) was added to a total mass of 2800.9 mg stool, and the mixture was left to soak for 10 minutes at room temperature. The stool mixture was homogenized by stirring and scraping stool solids against walls of the tube with a sterilized metal weighing spatula while swirling the tube and vortexing lightly at intervals until it reached a uniform consistency with only very small solid particles remaining. During homogenization, an additional 5ml of TE buffer was added and the resulting volume of the stool homogenate was 12.5ml. 250 aliquots of 50μl each were collected into individually pre-weighed Covaris Screw Cap AFA tubes (Covaris). The weights of samples in AFA tubes were recorded and stored at 4°C until ready for AFA or bead-beating lysis treatment (Figure 1A).

**Figure 1:**
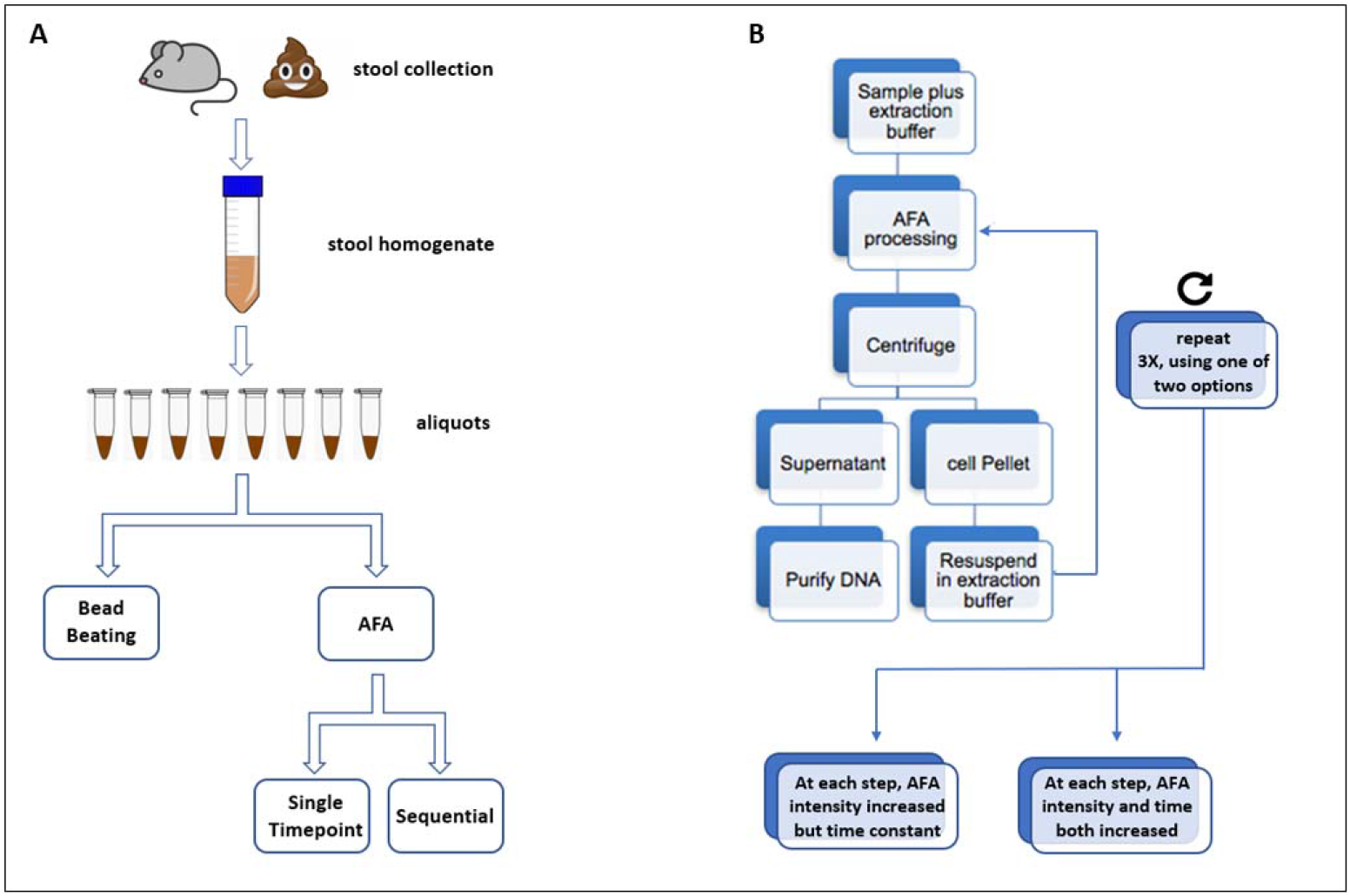
Experimental design and conditions Experimental design for balanced comparison of lysis methods. (A) Stool from healthy wild-type C57BL/6J mouse was collected, homogenized after addition of TE buffer and divided into identical aliquots. Groups of ten aliquots were processed using different bead-beating and AFA settings. AFA treatments included both single-timepoint and sequential lysis protocols. (B) Sequential AFA lysis protocol: Starting with a single aliquot, three progressively higher levels of AFA energy were applied. After each step, 450 μl of the lysate was reserved and replaced with equivalent volume of lysis buffer.

For AFA lysis, batches of 20 of the above identical samples were removed at a time from refrigeration, and 450μl of lysis buffer warmed to 60°C was added to each tube (Qiagen Solution MBL, Qiagen). Samples 1-250 were subjected to AFA treatment with the Covaris S220 system at the power levels and time points listed in Table 1. Samples 1-10 served as control treatment and did not undergo AFA processing. For sequential lysis treatment groups, after the initial AFA step, tubes were centrifuged for 10 minutes (3000 rcf at room temperature) and 450 μl supernatant was removed for DNA extraction and replaced with 450 μl of warmed lysis buffer.

**Table 1:** Detailed explanation of treatment groups for bead beating and AFA lysis

| Group | Explanation of lysis settings | AFA Joules |
| --- | --- | --- |
| NAC* | no-AFA negative control | 0 |
| BLP* | bead beating, low setting | NA |
| BMP | bead beating, medium setting | NA |
| BHP* | bead beating, high setting | NA |
| SL1* | single timepoint AFA, low setting, 1 min | 420 |
| SL2 | single timepoint AFA, low setting, 2 mins | 840 |
| SL4 | single timepoint AFA, low setting, 4 mins | 1680 |
| SM1 | single timepoint AFA, medium setting, 1 minute | 840 |
| SM2* | single timepoint AFA, medium setting, 2 mins | 1680 |
| SM4 | single timepoint AFA, medium setting, 4 mins | 3320 |
| SH1 | single timepoint AFA, high setting, 1 minute | 1440 |
| SH2 | single timepoint AFA, high setting, 2 mins | 2880 |
| SH4* | single timepoint AFA, high setting, 4 mins | 5760 |
| QL1 | sequential AFA, low setting first subaliquot, 1 minute | 420 |
| QL2 | sequential AFA, low setting second subaliquot, 2 mins | 840 |
| QL4 | sequential AFA, low setting final subaliquot, 3 mins | 2520 |
| QM1* | sequential AFA, medium setting first subaliquot, 1 minute | 840 |
| QM2* | sequential AFA, medium setting second subaliquot, 2 mins | 1680 |
| QM3* | sequential AFA, medium setting final subaliquot, 3 mins | 2520 |
| QH1 | sequential AFA, high setting first subaliquot, 1 minute | 1440 |
| QH2 | sequential AFA, high setting second subaliquot, 2 mins | 2880 |
| QH3 | sequential AFA, high setting final subaliquot, 3 mins | 4320 |
| CL1 | sequential AFA, constant time, varying intensity, low setting, 1 minute | 420 |
| CM2 | sequential AFA, constant time, varying intensity, medium setting, 2 mins | 1260 |
| CH3 | sequential AFA, constant time, varying intensity, high setting, 3 mins | 2700 |
| VL1 | sequential AFA, varying time, varying intensity, low setting, 1 minute | 420 |
| VM3 | sequential AFA, varying time, varying intensity, medium setting, 3 mins | 2100 |
| VH7* | sequential AFA, varying time, varying intensity, high setting, 7 mins | 7860 |
| CTII | sequential AFA metasample informatically merging CL1, CM2 and CH3 | NA |
| ITII | sequential AFA metasample informatically merging VL1, VM3 and VH7 | NA |

\* three samples from each of these groups also profiled by shotgun metagenome sequencing

The tube was then processed twice more with higher levels of AFA treatment, either with lysis time at each step held constant but AFA intensity increased (CTII treatment group; <u>C</u>onstant <u>T</u>ime <u>I</u>ncreasing <u>I</u>ntensity), or with lysis time and AFA intensity both increased (ITII treatment group; <u>I</u>ncreasing <u>T</u>ime <u>I</u>ncreasing <u>I</u>ntensity) (Table 1). Thus, each sequential lysis treatment produced three final samples, which were analyzed both as individual data points and also as a “metasample” using bioinformatically combined data from all three subsamples. AFA energy in joules transduced into the samples was calculated as the product of three instrument settings: acoustic energy in Watts (PIP), the duty factor (i.e., percent of lysis time that the transducer is on), and the duration (in seconds) of the AFA treatment. For the bead-beating lysis group (samples through 251-280), 0.05 mL of stool homogenate was aliquoted into a Qiagen PowerBead Plate (0.1 mm beads) and subjected to bead beating lysis treatment with a Retsch MM4000 bead mill, using either low-power treatment at 10 Hz, medium-power treatment at 20 Hz, or high-power treatment at 30 Hz (Table 1). After bead beating or AFA lysis, samples were processed according to the Qiagen PowerMag Microbiome DNA/RNA Isolation Kit protocols on an epMotion 5073 liquid handling robot (Eppendorf). Extracted DNA was quantified using the QIAxpert A260 dsDNA system (Qiagen) and plated on 96-well plates normalized to 100ng DNA per well using an epMotion 5073 liquid handler (Eppendorf).

### PCR amplification of 16S rRNA, shotgun metagenome library construction and Illumina sequencing

The 254bp V4 segment of the 16S ribosomal RNA subunit was amplified by a 17-cycle touchdown PCR using the following protocol: 12.5 μl Phusion 2X Master Mix (New England BioLabs),1 μl each of 16S V4 forward and reverse primers (primer pair 515F-806R), 100ng DNA in 10.5 μl elution buffer from previous step. PCR negative controls included lysis buffer only and PCR-grade water only, while positive controls were previously extracted fecal DNA and a bacterial mock community DNA. After PCR, amplicons were cleaned using Ampure beads (Beckman Coulter) using an epMotion 5075 liquid handling robot (Eppendorf) and 50 μl DNA was eluted with Qiagen EB buffer. For multiplexing, Nextera XT unique adapter indices (Illumina) were added to each sample in a 9-cycle PCR reaction prepared by adding 12.5 μl Phusion 2X Master Mix (NEB) and 2 μl each of Nextera XT S5 and N7 adapters. A second Ampure bead cleanup was performed on the epMotion 5075 using 80% ethanol, and amplicons were eluted in 25 μl of Qiagen EB buffer. DNA concentration was quantified using the KAPAQuant system (Roche) on the QuantStudio 6 Flex (Life Technologies). Next, an equimolar pooled library with 50ng of each amplicon was prepared using the epMotion 5075 system. The final pooled 16S amplicon library was diluted to 8pM, and after adding a 10% phiX spike-in, was loaded on the Illumina MiSeq system and sequenced using paired-end 300bp reads, designed to partially overlap at the middle of the 254bp V4 amplicon. For shotgun metagenomic sequencing, three samples each were randomly selected from 11 treatment groups representing the breadth of AFA and bead-beating treatments: NAC (non-AFA control), SL1 (single timepoint, low AFA power, 1 minute treatment time), QL1 (sequential treatment, low AFA, 1 minute), QL4 (sequential treatment, low AFA, 4 minutes), QH1 (sequential treatment, high AFA, 1 minute), QH3 (sequential treatment, high AFA, 3 minutes), CL1 (constant time, low AFA, 1 minute), VH7 (variable time, high AFA, 7 minutes), BLP (bead beating, low power), BMP (bead beating, medium power), SM4 (single timepoint, medium AFA, 4 minutes), and SH1 (single timepoint, high AFA, 1 minute). 10ng of the same DNA used for 16S amplicon sequencing was processed according to the Nextera Flex library construction protocol (Illumina) and automated on the Eppendorf epMotion 5075. Shotgun libraries were sequenced on the Illumina NextSeq system using 2×75bp paired-end reads.

### Bioinformatics analysis of 16S amplicon and shotgun metagenome sequencing data

Raw sequencer output files (in FASTQ format) were demultiplexed using bcl2fastq to generate paired-end FASTQ files for each sample. For 16S amplicon reads, these were pre-processed and analyzed using QIIME2 version 2-2018.2.^24^ Within QIIME2, the DADA2 algorithm was used for error modelling and filtering the raw data.^25^ Taxonomic classification was performed using the QIIME2 feature-classifier plugin trained on the Silva 132 database.^26^ Microbiome alpha and beta-diversity analyses were performed using the QIIME2 diversity plugin at a sampling depth of 30000. 294 (92.74%) of the samples were retained at this sampling depth. The Qiime 3D PCoA plots showing beta diversity differences were generated using Emperor.^27^ For analyzing shotgun metagenomic data, an in-house bioinformatics package in R was developed (JAMS version 1.19; https://github.com/johnmcculloch/JAMS_BW) which is described in a recent study.^28^ A detailed description of the shotgun metagenome analysis methods is also provided in the supplemental files for this study.

## Results

### Experimental design for comparison of bacterial lysis techniques

In this study, we present results from a balanced comparison of two microbiome lysis methods (bead-beating and AFA), with multiple treatment conditions for each method. For easier interpretation of the results below, we describe here the experimental design of our study (Figure 1). Using identical aliquots of homogenized mouse stool as a substrate (see Methods), bead-beating lysis was performed using three different instrument settings (low, medium and high, denoted BLP, BMP and BHP, respectively), while the AFA platform was tested across a wide range of settings with final energy delivered into the sample, ranging from 420-7860 joules. Further, our study included two types of AFA lysis treatment. The first was a single-timepoint approach, where individual aliquots were subjected to a single level of AFA energy. Separately, to account for variation in bacterial cell wall thicknesses, we designed sequential lysis protocols where each stool aliquot was subjected to three progressively higher AFA energy levels. In this approach, at each step, the sample was centrifuged, and a part of the supernatant was passed on to the next higher energy level, using variation in both lysis time as well as energy settings (see Methods). Ten technical replicates were used for all treatment groups; a detailed description of treatment conditions is provided in Table 1 and Figure 1. Technical process controls in our experimental design included both sample negative controls (lysis buffer only) and treatment negative controls (sample in lysis buffer but no bead beating or AFA treatment).

### DNA yields and integrity in bead-beating versus AFA lysis

As a primary technical comparison of bead beating and AFA (i.e., before generating NGS data), we compared the yields and integrity of DNA released by the two methods. We found that across all energy settings, AFA released significantly more DNA than bead-beating (two-tailed Student’s T-test, p<0.0001) (Figure 2A). Control samples placed in lysis buffer but not subjected to AFA treatment showed a similar yield to the medium and high-power bead beating groups, while the low-power bead beating group had the lowest yield among all. For the sequential AFA experiments, the second and third timepoints within each group yielded lesser DNA than the first (Figure 2B), which was expected, since sub-aliquots from the first lysis supernatant were used for the two subsequent timepoints. However, cumulative DNA output from the sequentially lysed AFA samples was substantially higher than both bead-beating and single-timepoint AFA treatments (two-tailed Student’s T-test, p<0.0001). In general, irrespective of single-timepoint or sequential treatment, our results point to an increase in lysis efficiency and correspondingly greater DNA yield for AFA as a microbiome sample extraction technique relative to bead beating.

**Figure 2:**
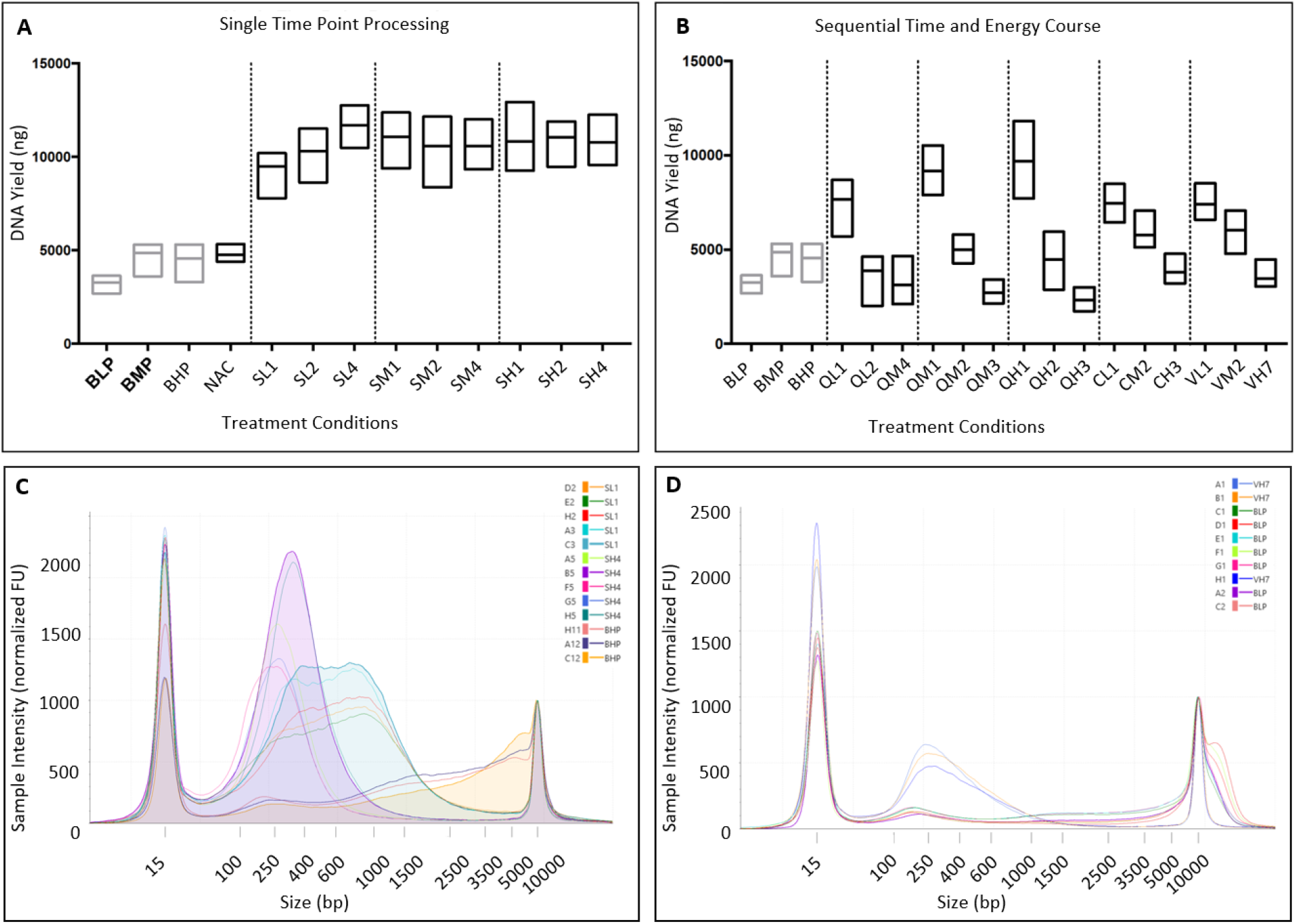
DNA yields and integrity from AFA and bead-beating protocols DNA yield and integrity from bead-beating and AFA protocols. (A) Single-timepoint AFA lysis compared with bead-beating: comparison of low, medium and high-power bead-beating groups (BLP, BMP and BHP, respectively) with low, medium and high-power AFA treatment for one, two and four minutes respectively (SL1/SL2/SL4; SM1/SM2/SM4; SH1/SH2/SH4). NAC samples were negative controls not treated with either bead-beating or AFA. Y-axis shows DNA yield in nanograms. (B) Sequential AFA lysis compared with bead-beating; details of AFA settings (groups QL1 through VH7) provided in Table 1. (C) Comparison of DNA integrity from AFA and bead-beating (four out of groups shown) (D) Underfragmented and overfragmented DNA from BLP and VH7 treatment (note shoulder at extreme right of electropherogram and peak <250bp, respectively).

Concurrently, we analyzed the relative integrity of DNA derived from bead-beating and AFA lysis. Most settings used for either technique resulted in NGS-compatible DNA fragment sizes (i.e., larger than the amplicon sizes commonly used for 16S analysis or the average insert sizes of a shotgun metagenome library; Figures 2C, four representative treatment groups shown for clarity). However, two experimental groups reveal shortcomings when extreme lysis settings are used. Bead beating at low settings (BLP) resulted in an abundance of DNA at very large fragment sizes (> 10000bp) (Figure 2D; note shoulder at extreme right of electropherogram). This suggests that incomplete lysis may be happening at that energy level, which is further supported by low alpha diversity in this group of samples (see below). At the other end of the lysis energy spectrum, samples in the highest AFA energy category (VH7) were fragmented to an average size of ~250bp (Figure 2D), which is smaller than optimal for both 16S amplicon sequencing and shotgun metagenomics. Again, this was reflected in the VH7 group having the lowest alpha diversity in our study (see below).

### Alpha diversity patterns in AFA and bead beating samples

For microbiome profiling of the stool homogenate, we amplified and sequenced the V4 hypervariable region of the 16S ribosomal subunit. We found that relative to all three bead-beating categories, single-timepoint AFA samples as a group showed a significantly higher alpha diversity (Figure 3A; pairwise Kruskal-Wallis test on observed OTU counts, all comparisons q-value< 0.05). Changes in alpha diversity among the three bead-beating energy settings were not pronounced, apart from a marginally significant difference between the BLP and BMP groups. Within the AFA energy setting spectrum, levels of alpha-diversity were highest in samples with lower energy lysis settings (Figure 3B). For example, two treatment groups lysed with 420 joules of AFA energy (CL1 and SL1) were both significantly higher in alpha diversity than the AFA groups with very high energy settings (VH7 and QH3, 7860 and 4320 joules of energy, respectively) (pairwise Kruskal-Wallis tests, q<0.00001 for both comparisons). In other AFA groups with intermediate settings, observed levels of alpha diversity fell between these extremes. Interestingly, single-timepoint treatment groups processed with high amounts of AFA energy (e.g. SH4, processed with 5760 joules) did not have decreased alpha diversity. Hence, the observed drop in alpha diversity in VH7 and QH3 likely reflects over-fragmentation of DNA when high AFA energy is applied to sub-aliquots of a sequentially lysed sample that have already been though lower levels of AFA treatment, which is also supported by DNA integrity as seen in the electropherograms of Figure 2C.

**Figure 3:**
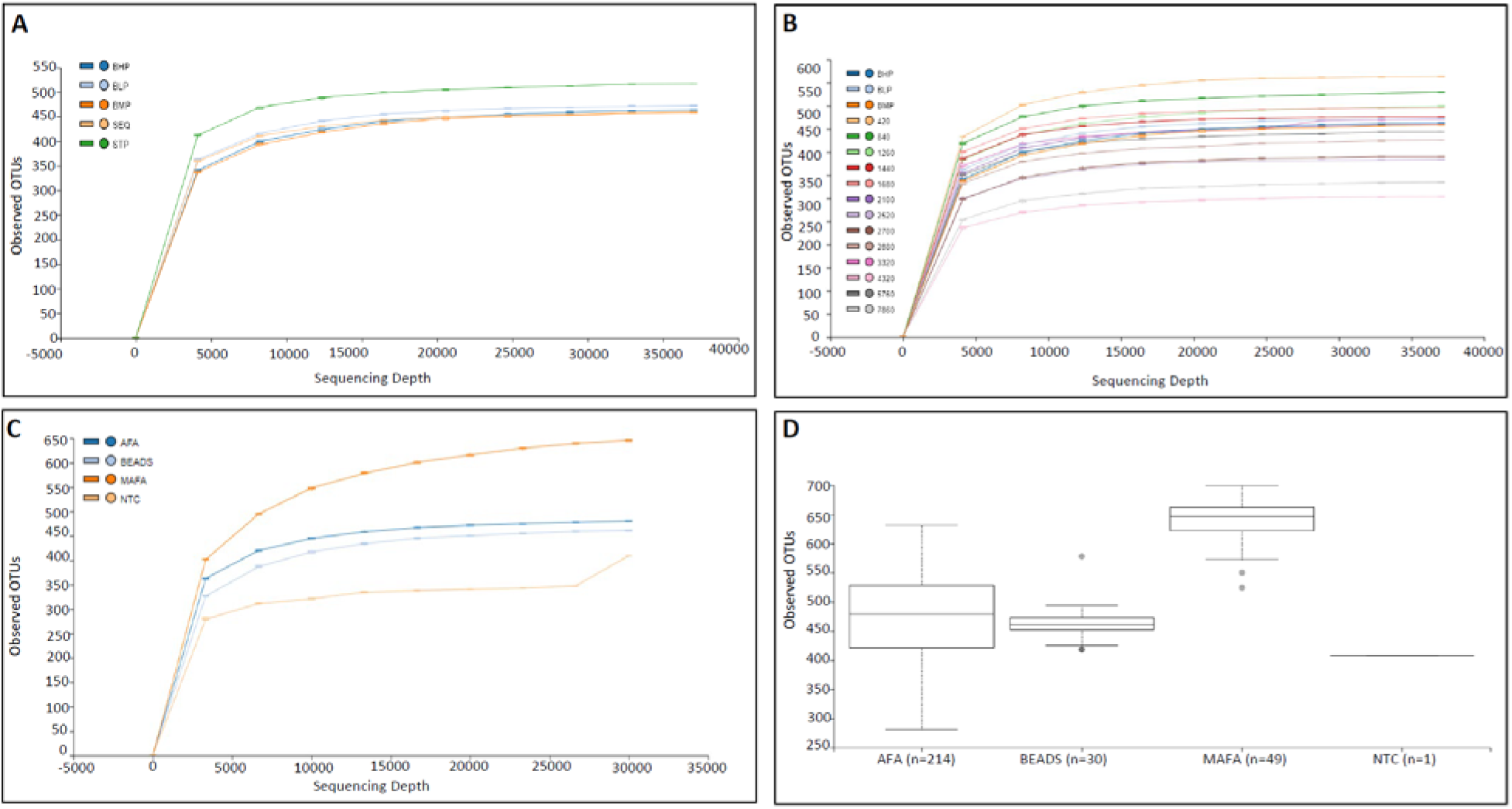
Alpha diversity differences in AFA and bead-beating samples Alpha diversity differences between bead-beating and AFA samples. (A) Comparison of single-timepoint and sequential AFA samples (STP and SEQ, respectively) with low, medium and high-power bead-beating groups (BLP, BMP and BHP, respectively). (B) Alpha diversity in AFA treatment groups with different amounts of acoustic energy transduced into the sample aliquot; note higher levels of diversity in low-energy AFA groups (420 and 840 joules) relative to highest energy AFA treatment (5760 and 7860 joules, respectively). (C) Analyses of AFA metasamples (MAFA) comprising three sub-aliquots from a single sequential lysis treatment analyzed in aggregate, showing substantially higher alpha diversity relative to bead-beating (BEADS) and single-timepoint samples (AFA). NTC: no-template control; (D) Statistical comparison of alpha diversity in metasamples with single-timepoint AFA treatment and bead-beating (pairwise Kruskal-Wallis tests on observed OTU counts, all comparisons q-value <0.05). For all panels; Y-axes show alpha diversity measured as observed OTUs; For panels A-C, X-axes show sequencing read depth; for panel D, x-axes show number of samples in each category.

Next, we examined the additive benefits of our sequential AFA lysis protocol over single-timepoint lysis. To produce a data set that was representative of the cumulative DNA content at the end of sequential lysis, we combined all the NGS reads from all three sub-aliquots of the original stool homogenate tube (creating what we refer to hereafter as “metasamples”). Next, we bioinformatically down-sampled these metasamples to match the depth of sequencing in groups that were processed in a single timepoint (see Methods). Comparison of metasamples with both single-timepoint AFA treatment and bead-beating revealed a particularly large and significant increase in the number of observed OTUs (Figure 3C-D; pairwise Kruskal-Wallis tests on observed OTU counts, all comparisons q-value <0.0001). Specifically, this particular result highlights the possibility that in existing one-step microbiome lysis protocols, an unknown number of bacterial taxa may escape detection altogether, and that sequential lysis accounting for differences in cell wall rigidity captures greater diversity in microbiome samples.

### Beta diversity and reproducibility patterns in AFA relative to bead beating

Moving beyond the observed differences in alpha diversity, we analyzed between-group or beta diversity in AFA and bead-beating samples (using the weighted UniFrac distance metric). This revealed that AFA energy (in Joules) delivered into the sample was the greatest contributor to beta diversity (Figure 4A). Using Principal Coordinates Analysis (PCoA), we determined that this factor accounted for 86.07% of the total variation (located along the first PCoA axis).

**Figure 4:**
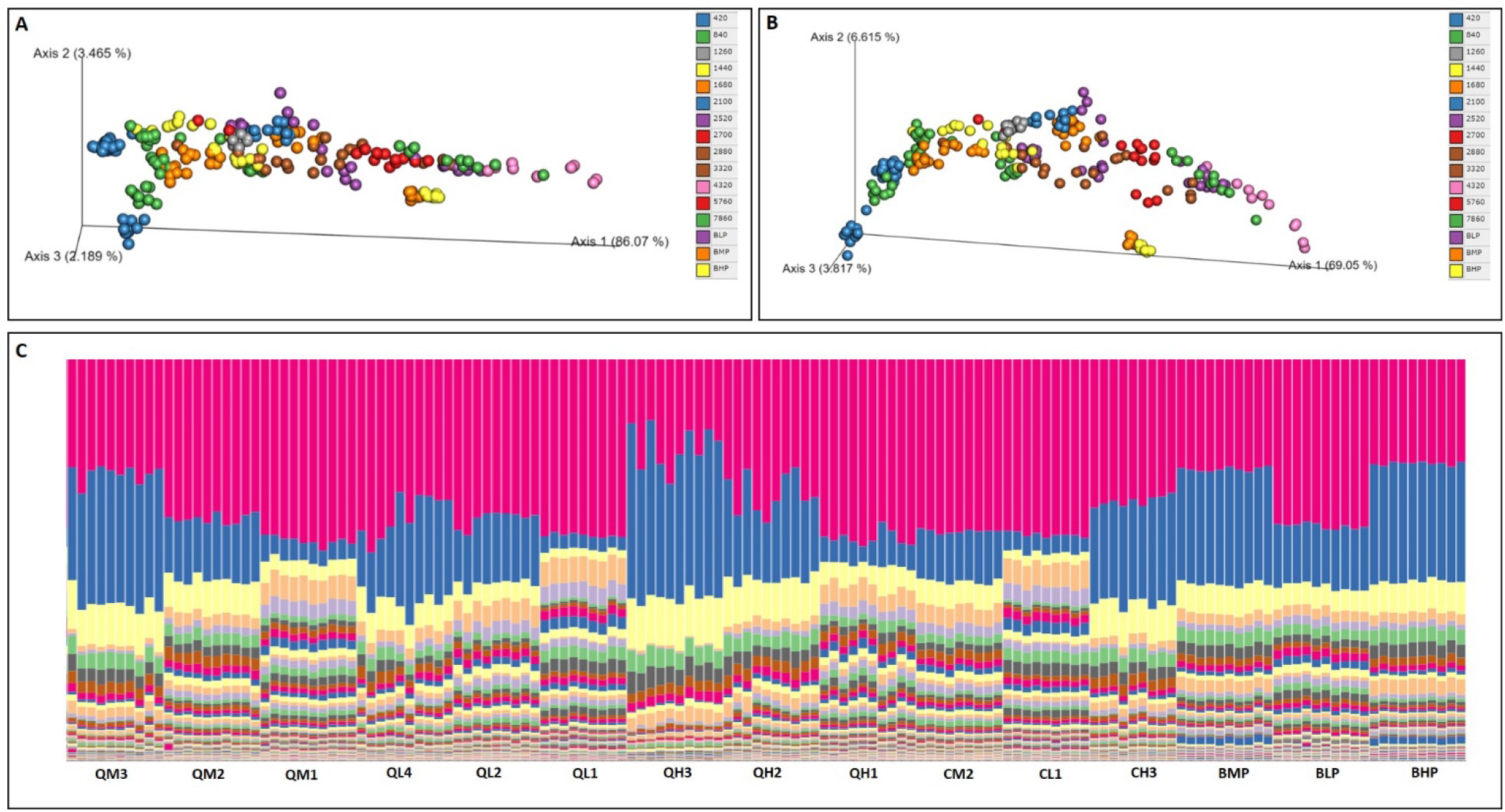
Beta diversity differences in AFA and bead-beating samples Beta diversity differences between bead-beating and AFA samples. (A) Principal Coordinates Analysis (PCoA) of weighted UniFrac distances between different AFA treatment groups (clustered by AFA energy in joules) and bead-beating groups. Note linear clustering of different AFA sample groups in order of increasing AFA energy from the origin outwards along Axis 1 (86.07% of total variation). Medium and high-power bead-beating groups (BMP and BHP) occupy an intermediate position. (B) Alternative indices of beta diversity (Bray-Curtis dissimilarity) show similar results to weighted UniFrac. (C) Taxonomic diversity (assessed as stacked taxa bar plots) in sets of ten technical replicates for AFA and bead-beating treatment groups. Given that all aliquots were identical, taxonomical pattern differences clearly separating each group of ten replicates shows effects of lysis method and setting on beta diversity results.

Specifically, we found that different AFA sample groups clustered linearly (from the origin outwards) in order of increasing AFA energy, with the medium and high-power bead-beating groups (BMP and BHP) occupying an intermediate position. Alternative indices of beta diversity (such as the Bray-Curtis dissimilarity) showed similar results to the above findings (Figure 4B). Beyond taxonomy-agnostic beta-diversity analyses using PCoA, we also analyzed taxonomic differences among the AFA and bead-beating lysis groups. Since identical stool homogenate aliquots were used for all treatment groups and all samples were processed uniformly post-lysis, in our experimental design, differences in species composition can be directly attributed to lysis technique. At the genus level (corresponding to Qiime2 Level 6), stacked bar plots of taxonomic abundances showed both consistency within the ten replicates used for each treatment group, and a clear pattern of between-group differences (Figure 4C).

To identify taxonomic patterns linked to lysis technique, we used linear discriminant analysis effect size analysis (LefSe) to compare genus-level differences between bead-beating and AFA treatment.^29^ This revealed a consistent trend for AFA treatment groups to detect a higher number of taxa with increased efficiency relative to bead-beating treatment groups (Figure 5A).

**Figure 5:**
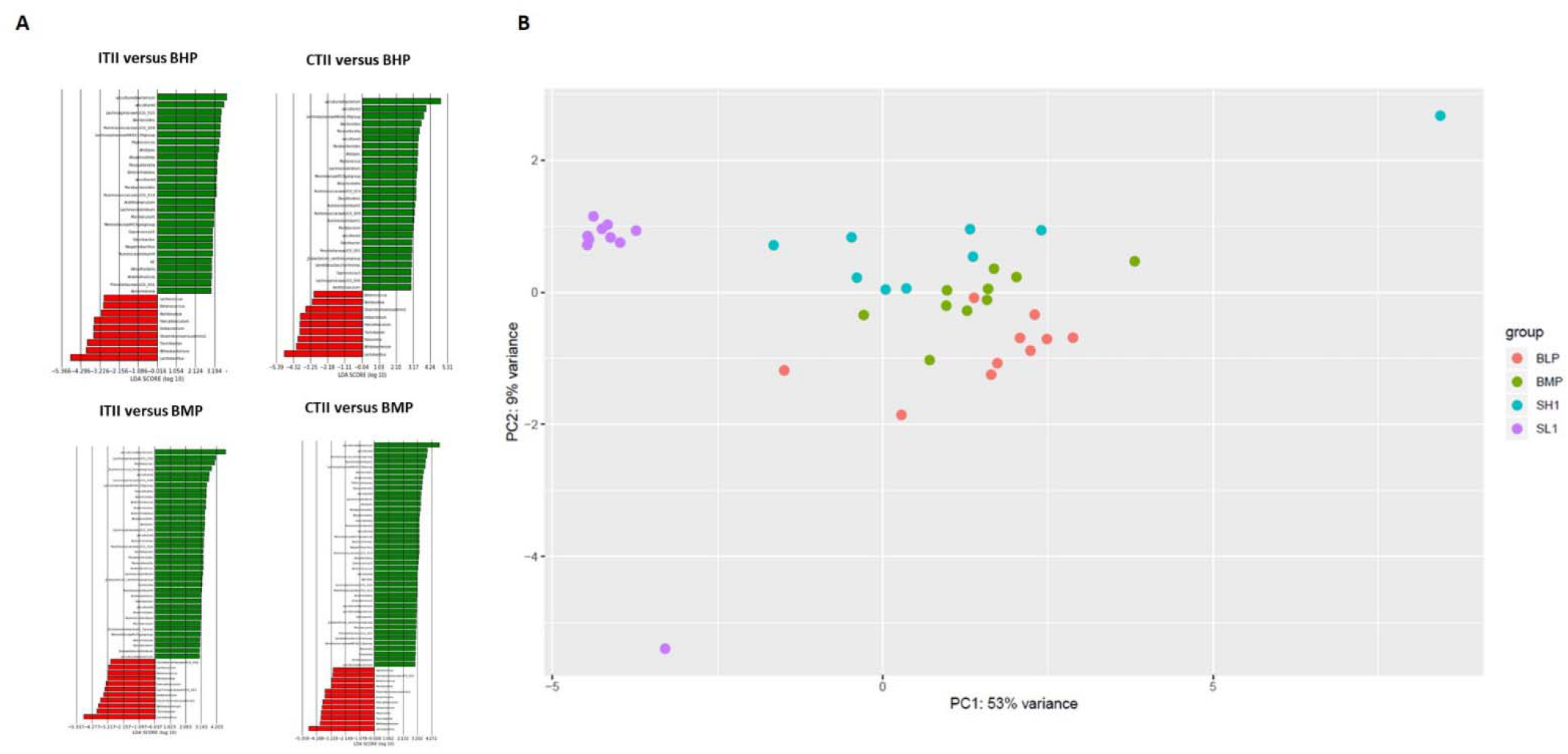
Beta diversity and reproducibility analysis of AFA and bead-beating samples Analyses of differentially detected taxa in bead-beating and AFA and reproducibility assessment. (A) LEfSe (Linear discriminant analysis Effect Size) results from comparison of medium and high-power bead-beating groups (BMP and BHP, respectively) with metasamples from sequential AFA lysis treatment (CTII; <u>C</u>onstant <u>T</u>ime <u>I</u>ncreasing <u>I</u>ntensity and ITII; <u>I</u>ncreasing <u>T</u>ime <u>I</u>ncreasing <u>I</u>ntensity, respectively). Green and red bars represent taxa that were found with increased efficiency in AFA and bead-beating lysis, respectively. (B) Principal Components Analysis (PCoA) results of clustering in groups of ten technical replicates from low and medium power bead-beating (BLP and BMP, respectively) compared with mild and extreme AFA settings (SL1 and VH7, respectively). Note substantially tighter clustering in SL1 group relative to all others.

This result requires careful interpretation. Since identical stool homogenate aliquots were used in all cases (see Methods), under the null assumption that AFA and bead-beating methods are equally efficient for cell wall lysis, LefSe analysis would return no significant differences, which was not the case in our results. Given the presence of taxonomic differences returned from the same starting material, the next testable hypothesis would be that both techniques will return equal numbers of species detected at higher levels relative to each other, which again is not the case. Hence, the observed pattern (higher number of taxa with higher representation in AFA) reflects the ability of AFA to capture more DNA from more species in the same starting bacterial community relative to bead-beating. Moving beyond diversity analyses, although our study was not originally designed to test reproducibility, we investigated this aspect of differences between lysis techniques, since all the treatment groups for both AFA and bead-beating included ten technical replicates. Focusing on the low and medium power bead-beating groups (BLP and BMP, respectively) and a subset of the AFA groups chosen to represent different levels of acoustic energy (SL1, SH1 and VH7), we first used principal components analysis (PCA) to study dispersion within each group of ten technical replicates. We found that samples in the SL1 group clustered much more closely than any of the other groups (Figure 5B), suggesting that low-energy AFA lysis may be a way to improve reproducibility in microbiome data over bead-beating protocols.

### Species-level analysis of differences between bead-beating and AFA lysis

As a follow-up to the 16S amplicon sequencing experiments, we chose a subset of the AFA and bead-beating samples for further analysis using shotgun metagenome sequencing (Table 1). After reference-agnostic metagenome assembly (see Methods), non-supervised hierarchical clustering analysis of groups from two extremes of the AFA treatment spectrum (SL1 and VH7) compared with the BMP group showed that SL1 samples were tightly clustered relative to the other groups (Figure 6A). By annotating species differences between the three groups with either Enzyme Classification numbers^30^ or Pfam IDs,^31^ we found that lysis-induced noise in microbiome studies is reflected not just in taxonomic composition but also in functional interpretation of beta diversity results (Figure 6 B,C). Finally, although the lysis substrate we selected for our experiments (mouse stool homogenate) is expected to contain very few eukaryotic species, we extended our shotgun metagenome analysis beyond the prokaryotic microbiome to test whether AFA versus bead-beating lysis differentially affected recovery of eukaryotic DNA. This revealed that AFA samples as a group had higher levels of eukaryotic DNA (Figure 6D). When eukaryotic DNA content was further analyzed at the treatment subgroup level, this observed increase in both the AFA groups (SL1 and VH7), relative to the BMP group. Possibly in keeping with the 16S amplicon sequencing results, the SL1 AFA group had the highest relative `abundance` (Figure 6E). The lone fungal species detected in these analyses (*Phycomyces blakesleeanus*, whose cell wall is largely composed of tough chitin fibrils) was significantly higher in SL1 samples, as was the protozoan *Trichomonas vaginalis*, which is known to form hard cyst-like structures in the course of its lifecycle (Supplemental Figure 1). We suggest that for samples with higher fungal biodiversity or known to contain lysis-resistant species, AFA deserves consideration as an alternative lysis technique.

**Figure 6:**
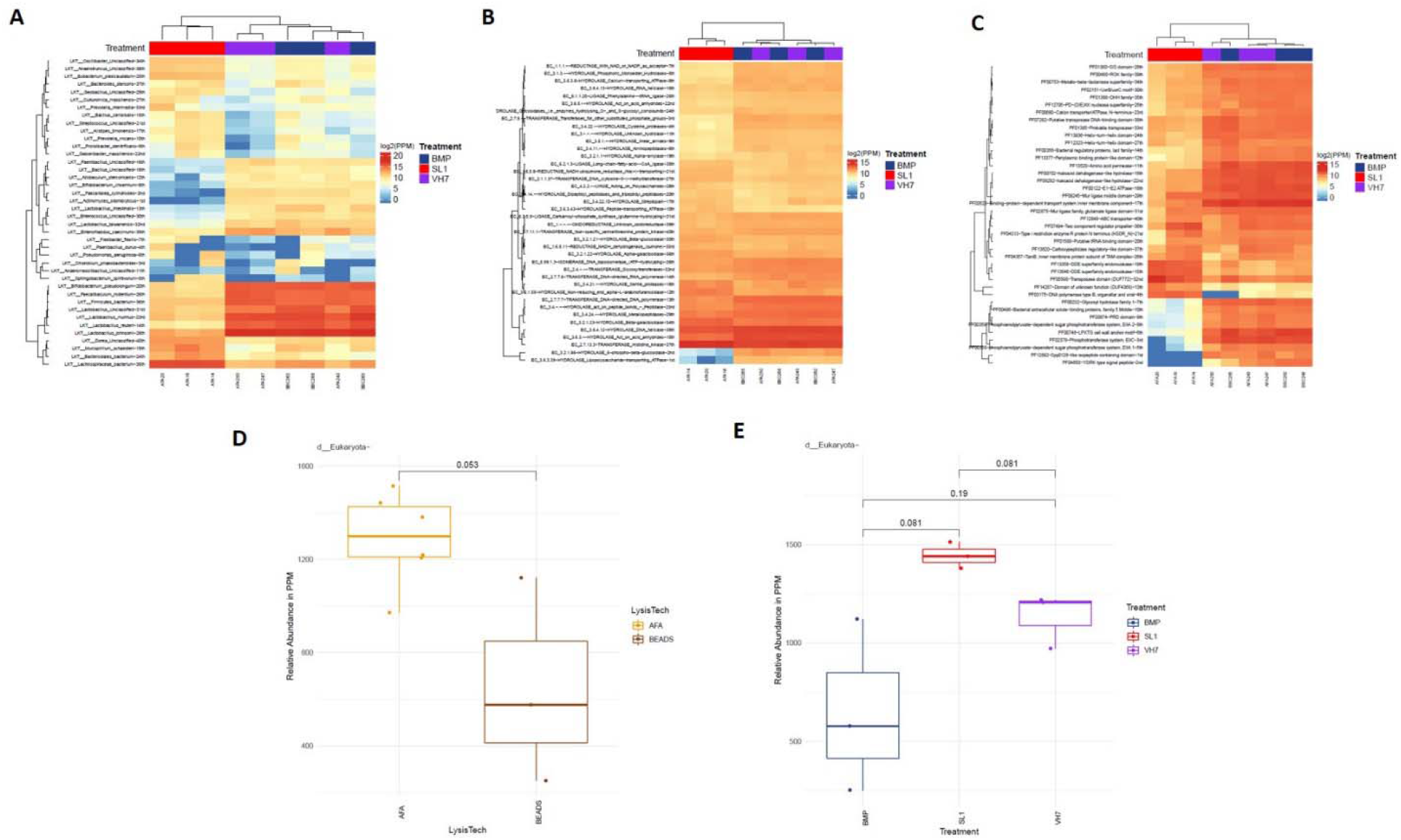
Shotgun metagenome sequencing Shotgun metagenomics results from bead-beating and AFA samples. (A) Unsupervised hierarchical clustering of species-level differences between medium-power bead-beating group (BMP) and two AFA groups with gentle and harsh settings (SL1 and VH7, respectively). (B) Enzyme Commission number analysis showing bacterial enzymes with non-biological (i.e. lysis-induced) differences between SL1, BMP and VH7 (C) Pfam analysis showing protein families with non-biological (i.e. lysis-induced) differences between SL1, BMP and VH7 (D) Relative abundance in parts per million (PPM, Y-axis) of eukaryotic DNA between AFA and bead-beating samples (E) Relative abundance in parts per million (PPM, Y-axis) of eukaryotic DNA between BMP, SL1 and VH7 groups.

## Discussion

Recent years have seen a rapid surge in our knowledge of how the microbiome modulates human health and disease.^32^ However, the technical aspects of microbiome sequencing are still an evolving field.^6, 7^ In this context, sources of noise in microbiome studies^33^ need to be much better understood (along with the ways in which they affect reproducibility of results).^4, 5^ Here, we revisited cell wall lysis as a fundamental aspect of microbiome sequencing protocols.^34^ We objectively compared existing lysis protocols with AFA, which is widely used in other NGS domains but has not been explored as a microbiome tool.^20–22^ AFA technology has characteristics that led us to consider it as a logical choice for processing microbial samples, such as precise scalability across a broad range of energy levels, isothermal lysis conditions and the ability to process especially lysis-resistant bacterial species by transducing large amounts of acoustic energy. Using identical aliquots of a mouse stool homogenate as the substrate, we found that a one-step AFA protocol yields more DNA and uncovers greater microbiome diversity relative to bead beating. Further, a sequential AFA lysis protocol resulted in even more bacterial taxa being detected in NGS data, highlighting the possibility that current lysis methods may miss components of microbiome diversity. The objective of our work was not to find a single AFA-based protocol that is optimal for microbiome sequencing. Indeed, the tremendous structural diversity of bacteria^23, 35^ makes it likely that any one-size-fits-all lysis setting (whether for AFA or another method) could induce a bias in the final NGS data resulting from differential lysis of taxa with based on their cell wall thicknesses. Rather, we suggest that the additional diversity found using AFA as a technology merits increased attention to lysis as a component of microbiome sequencing protocols. Relative to single-timepoint AFA, sequential lysis treatment with a combination of three progressively higher AFA settings uncovered even greater diversity of the native microbiome; however, the additional hands-on time needed for sequential lysis may prevent this from being a routinely used method. Although our study does not comprehensively address the question of reproducibility in microbiome protocols, we show evidence that AFA lysis resulted tighter clustering among groups of technical replicates, relative to bead-beating. Regarding functional differences in the microbiome, since we used identical stool homogenate aliquots for all experiments, the null expectation would be an absence of any significant differences between AFA and bead-beating treatment groups. In this context, we find that lysis-induced noise distorts the view of dysregulated functional mechanisms in microbiome analyses. Moving beyond prokaryotes, we found suggestive evidence that lysis methods affect the ability to capture eukaryotic components of the microbiome sample, particularly in the case of fungi.

Given that microbiome sequencing is an evolving technical field, our study highlights the need for careful consideration of cell wall lysis technique to bring the full diversity of a microbiome sample into the NGS read sampling space. Both from a basic and clinical research viewpoint, microbiome science encompasses an especially vast variety of biosample types. In this context, the scalable nature of AFA presents has the advantage of allowing lysis energy levels to be customized to the sample type. In a broader context, microbial “dark matter”, currently escaping detection in NGS-based profiling due to incomplete lysis, may be obscuring part of the biology in microbiome-based studies. As NGS-based microbiome applications move from basic research into the clinical realm, this problem will become increasingly relevant, particularly in the identification of pathological or causative organisms. In general, we suggest that the greater diversity achievable through AFA lysis may help in shedding some light on the “unknown” aspects of microbiome science.^36^ In a non-microbiome context, AFA instruments are already in extremely widespread use for DNA shearing in most laboratories utilizing the Illumina NGS platform; hence, AFA technology for lysis may already be available to a large section of the microbiome sequencing community.

For balanced consideration of our results, it is important to acknowledge some limitations both of AFA as a technique and of our current study. Without careful optimization for the specific starting material type, higher AFA settings may lead to DNA overfragmentation and subsequent loss of taxonomic diversity, highlighting the need for pilot experiments with this technique while developing lysis protocols. On the other hand, high-energy AFA lysis could be a valuable technique for processing particularly lysis-resistant microbiome sample types (e.g. fungal samples or endospore-forming bacteria).^37^ We suggest that our experimental design, which spans a very wide range of AFA settings up to the highest energy limits supported by the instrument, provide a spectrum of possible lysis protocols ranging from mild to extreme. Regarding lysis methods beyond AFA and bead-beating, we purposefully chose to exclude enzymatic lysis from this study, as we chose to focus on non-biological methods without concerns about lower general applicability to all target organisms irrespective of biochemical structure of the bacterial cell wall. However, it is conceivable that a combination of gentle AFA treatment and enzymatic lysis may be optimal for some microbiome sample types. Finally, for laboratories not routinely engaged in NGS protocols, the AFA instrument may be relatively inaccessible.

In conclusion, through this study, we show a promising new lysis method for microbiome science, and demonstrate through balanced comparison with existing methods, that it uncovers increased DNA yield and taxonomic diversity when applied to the same starting material. We hope that our results will lead to an increased focus on lysis as a source of noise in microbiome science. We suggest that the AFA platform, already in widespread use for nucleic acid fragmentation, offers potential for improving current protocols for microbial cell wall lysis.

## Acknowledgements

The authors would like to thank Giorgio Trinchieri and Colm O’hUigin for scientific guidance, John McCulloch for helping with computational analysis of metagenomics data, and Mary Pingitore for assistance with AFA technology. This project has been funded in whole or in part with federal funds from the Frederick National Laboratory for Cancer Research, National Institutes of Health, under contract HHSN261200800001E. The content of this publication does not necessarily reflect the views or policies of the Department of Health and Human Services, nor does mention of trade names, commercial products or organizations imply endorsement by the US Government. This research was supported in part by the Intramural Research Program of NIH, Frederick National Lab, Center for Cancer Research. NCI-Frederick is accredited by AAALAC International and follows the Public Health Service Policy for the Care and Use of Laboratory Animals. Animal care was provided in accordance with the procedures outlined in the “Guide for Care and Use of Laboratory Animals (National Research Council; 1996; National Academy Press; Washington, DC). SKS was an employee of Leidos Biomedical Research, Inc. at the time this research was conducted, but is currently employed by Kelly Government Solutions, Inc.

## Author contributions

GW, HK and SKS performed all experiments. All authors analyzed and interpreted data. SKS wrote the manuscript. All authors read and approved the final manuscript.

## Availability of data and materials

Sequencing datasets generated and/or analyzed during this study are available in the [NAME] repository, [PERSISTENT WEB LINK TO DATASETS] under accession number [SRA ACCESSION NUMBER]

