## Supplemental Methods and Figures for "Hidden components of microbiome diversity revealed by cell wall lysis with Adaptive Focused Acoustics"

Detailed description of shotgun metagenome analysis pipeline

For analyzing shotgun metagenomic data, we used a de novo bacterial genome assembly-based computational framework. Non-human reads were first assembled into contigs using MEGAHIT, typically yielding 50-200 contigs per bacterial species. After de novo assembly, all the contigs within a sample were classified using Kraken which counts the occurrence of words of size 31bp occurring in the contig to be classified. We previously built a 31-mer word spectrum database for all bacterial species in GenBank, plus all fungal, archaeal, protozoan and viral (along with the human and mouse reference genomes). Importantly, if the contig was from an unknown species, it was possible to classify it by the last common ancestor of all known species in our database which contain that k-mer spectrum. Therefore, a contig from a species or strain absent in GenBank was classified as an unknown species but identified as belonging to a certain taxonomical clade above species, such as genus, family, order, class or phylum, (denoted by the tags g__, f__, o__, c__ or p__, respectively) next to the taxonomic identification. A hierarchical combination of these tags was used to assign a “Last Known Taxon" or LKT to such contigs. Once contigs were classified into species or LKTs in this manner, the proteome they code for was predicted ab initio, by searching for non-overlapping open reading frames between probable start and stop codons with a Hidden Markov Model-based tool, thus enabling the assignment of proteins to individual taxa. The function of each protein thus detected was predicted using InterProScan. This enables the study of predicted proteomes from each metagenome by several different aspects, such as Pfams (protein families), EC numbers (Enzyme Commission numbers) and Gene Ontology (GO) terms. After taxonomic and functional qualitative information were determined, the "dose" of each contig, taxonomical clade and function was obtained by aligning the sequencing reads back to the contigs and deriving the relative abundance of each contig (and therefore, of each taxon and ultimately each function). Crucially, for calculating taxonomic relative abundance, reads which were NOT assembled into contigs (ie. the ones which did not align to contigs) are also retained. These unassembled reads were themselves identified by k-mer spectrum like the contigs, and their relative dose was added to the computation of taxonomic relative abundance within each sample. Thus, the taxonomic relative abundance was calculated considering 100% of all DNA bases sequenced for a given sample. Finally, all the data was integrated using a custom-built R package which plots heatmaps, correlation plots, ordination plots and the relative abundances of features of interest. Software used for the above data processing steps is available at <https://github.com/johnmcculloch/JAMS_BW>.

Supplemental Figure 1

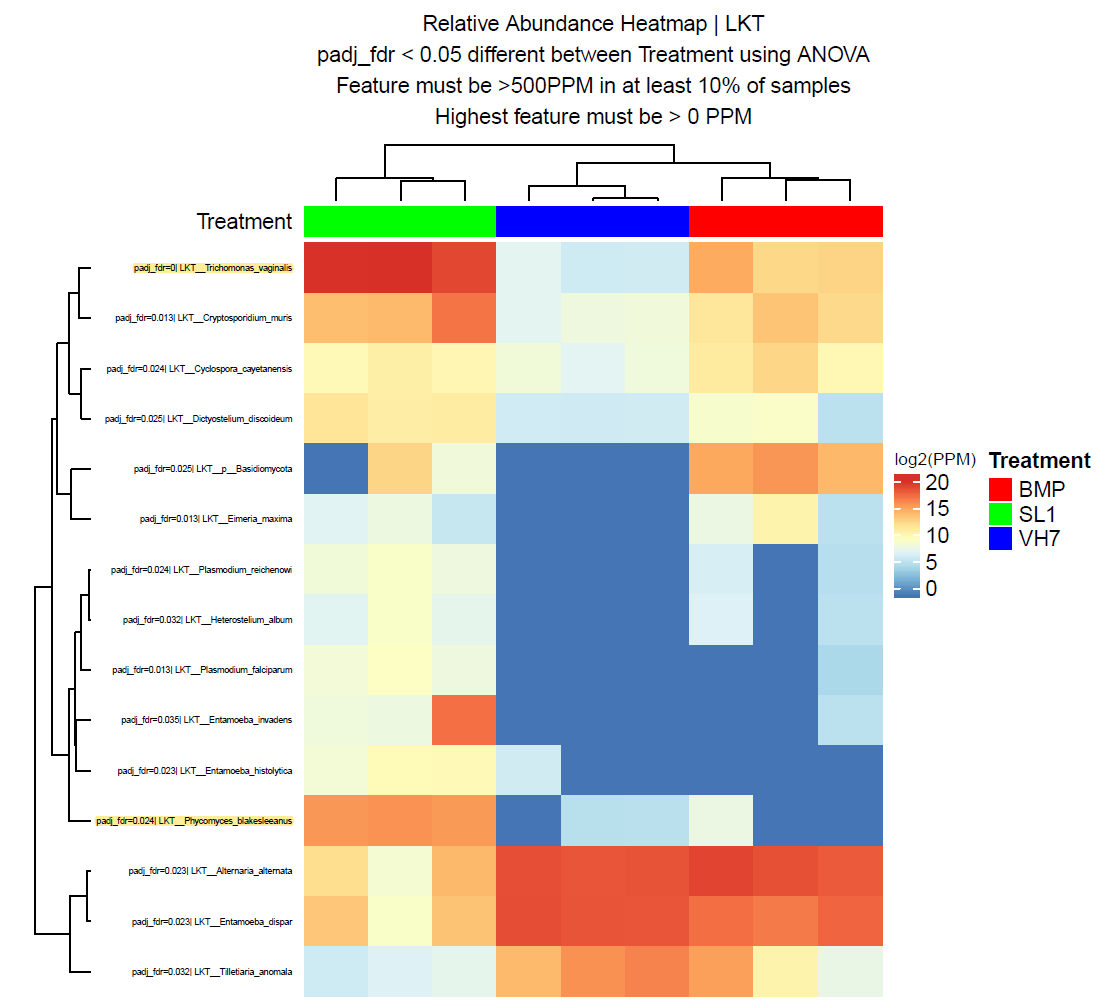

Figure legend: Supervised Clustering of shotgun metagenome sequencing results showing eukaryotic species differentially detected in AFA (SL1 and VH7) and bead-beating (BMP). Trichomonas vaginalis and Phycomyces blakesleeanus highlighted in yellow (top and bottom, respectively)
